# Multi-Modal Data Analysis for Alzheimer’s Disease Diagnosis: An Ensemble Model Using Imagery and Genetic Features

**DOI:** 10.1101/2021.05.07.443184

**Authors:** Qi Ying, Xin Xing, Liangliang Liu, Ai-Ling Lin, Nathan Jacobs, Gongbo Liang

## Abstract

Alzheimer’s disease (AD) is a devastating neurological disorder primarily affecting the elderly. An estimated 6.2 million Americans age 65 and older are suffering from Alzheimer’s dementia today. Brain magnetic resonance imaging (MRI) is widely used for the clinical diagnosis of AD. In the meanwhile, medical researchers have identified 40 risk locus using single-nucleotide polymorphisms (SNPs) information from Genome-wide association study (GWAS) in the past decades. However, existing studies usually treat MRI and GWAS separately. For instance, convolutional neural networks are often trained using MRI for AD diagnosis. GWAS and SNPs are frequently used to identify genomic traits. In this study, we propose a multi-modal AD diagnosis neural network that uses both MRIs and SNPs. The proposed method demonstrates a novel way to use GWAS findings by directly including SNPs in predictive models. We test the proposed methods on the Alzheimer’s Disease Neuroimaging Initiative dataset. The evaluation results show that the proposed method improves the model performance on AD diagnosis and achieves 93.5% AUC and 96.1% AP, respectively, when patients have both MRI and SNP data. We believe this work brings exciting new insights to GWAS applications and sheds light on future research directions.

## I. Introduction

Alzheimer’s disease (AD) is a chronic neurodegenerative disease, which will cause progressive damage to patients’ ability in memory, language, behavior, and problem solving [1]. An estimated 6.2 million Americans age 65 and older are suffering from Alzheimer’s dementia today. In 2019, official death certificates recorded 121,499 deaths from AD in the US, and this trajectory of deaths from AD was likely exacerbated in 2020 by the COVID-19 pandemic [2]. However, no treatment available at this time can cure or completely stop the progression of AD [3]. The difficulty in finding treatments for AD is most likely due to a lack of clear understanding of the cause and the fact that AD patients cannot be easily identified at early stages [4]. For this reason, early diagnosis is crucial for AD treatment and potential drug development.

Magnetic resonance imaging (MRI) is a widely used neuroimaging technology for AD diagnosis. An MRI is a pseud-3D image composed of 2D imaging slices. The voxels in MRIs are corresponding to the physical locations in patients’ brains. In recent years, machine learning techniques, especially convolutional neural networks (CNN), have been successfully applied in the diagnosis of AD patients using MRI images [5]–[10]. Compared with traditional computer-aided diagnosis tools, CNN does not rely on pre-defined, hand-crafted features and has shown promising results in medical imaging analysis tasks [11]–[16].

When making an AD diagnosis, CNN models normally take an MRI as input and output the probability that the patient has AD. The range of the probability is between 0 and 1, with 0 indicates a cognitively normal (CN) patient and 1 indicates an AD patient. The number of the predicted probability also indicates the degree of uncertainty. For instance, a probability of 0.05 may indicate `more than likely to be a CN patient`, 0.75 may indicate `probably is an AD patient`, and 0.5 may indicate `not sure`. Normally, when a CNN model makes decisions, 0.5 is usually used as the decision cut-off. For images with probability smaller than 0.5, the network predicts CN; otherwise, the network predicts AD. Such a method may be problematic when the predicted probability is close to 0.5, which indicates extreme uncertainty for the prediction (i.e., the model has low confidence in its prediction). In clinical practice, additional tests are ordered when a medical expert is not certain in his/her decision. Inspired by the clinical practice, we proposed to use additional information to refine the CNN model prediction when the predicted probability is extremely uncertain or prediction confidence is low.

Genome-wide association study (GWAS) is an approach used in genetics research to associate specific genetic variations with particular diseases by investigating associations between single-nucleotide polymorphisms (SNPs) and a variety of phenotypes [17], [18]. Genetic factors play an important part in the development of AD. Ridge et al. indicated that although most AD cases involve multiple genetic, environmental, and lifestyle factors, genetics can account for up to 53% of total phenotypic variance [19]. In the past decades, multiple GWAS studies have identified hundreds of SNPs associated with AD [20]. By using genetic linkage analysis, these SNPs can help to locate predictive AD biomarkers and genetic risk factors. Therefore, SNPs became a natural choice for the additional information to refine the CNN model prediction.

This study proposes to use SNPs to refine CNN model performance by ensemble the SNP predicting result with the CNN result when the prediction confidence of the CNN model is low. We evaluate the proposed method on the Alzheimer’s Disease Neuroimaging Initiative (ADNI) dataset (https://adni.loni.usc.edu). Our experimental result shows that the ensemble model using both SNP and MRI features improves the AD diagnosis performance than only using MRI images.

We consider our contributions to this work as the following:

- propose a cross-domain deep neural network that uses SNPs to refine CNN model prediction when the predicted confidence is low;
- demonstrate a novel way to use GWAS results in clinical practice by directly using SNPs in predictive models for AD diagnosis;
- improve the AD diagnosis performance from 92.4% AUC to 93.5% when patients have both MRI and SNPs;
- discuss the proposed method in detail and provide a clear research direction to future researchers.

## II. Approach

The proposed ensemble model contains two sub-networks, an image processing network and an SNP processing network (Figure 1). The image processing network and SNP processing network take an MRI and a set of SNPs as input and predict whether the patient is AD or CN separately. An ensemble gate (*σ*) is included in the ensemble model. If the image processing network has low confidence, the ensemble gate is open, and the SNP prediction is used to refine the image prediction. Otherwise, the ensemble gate remains closed; the image prediction result is used as the final prediction for the patient. Both of the networks can be trained jointly or separately. In this project, we train the networks separately for simplicity.

**Fig. 1:**
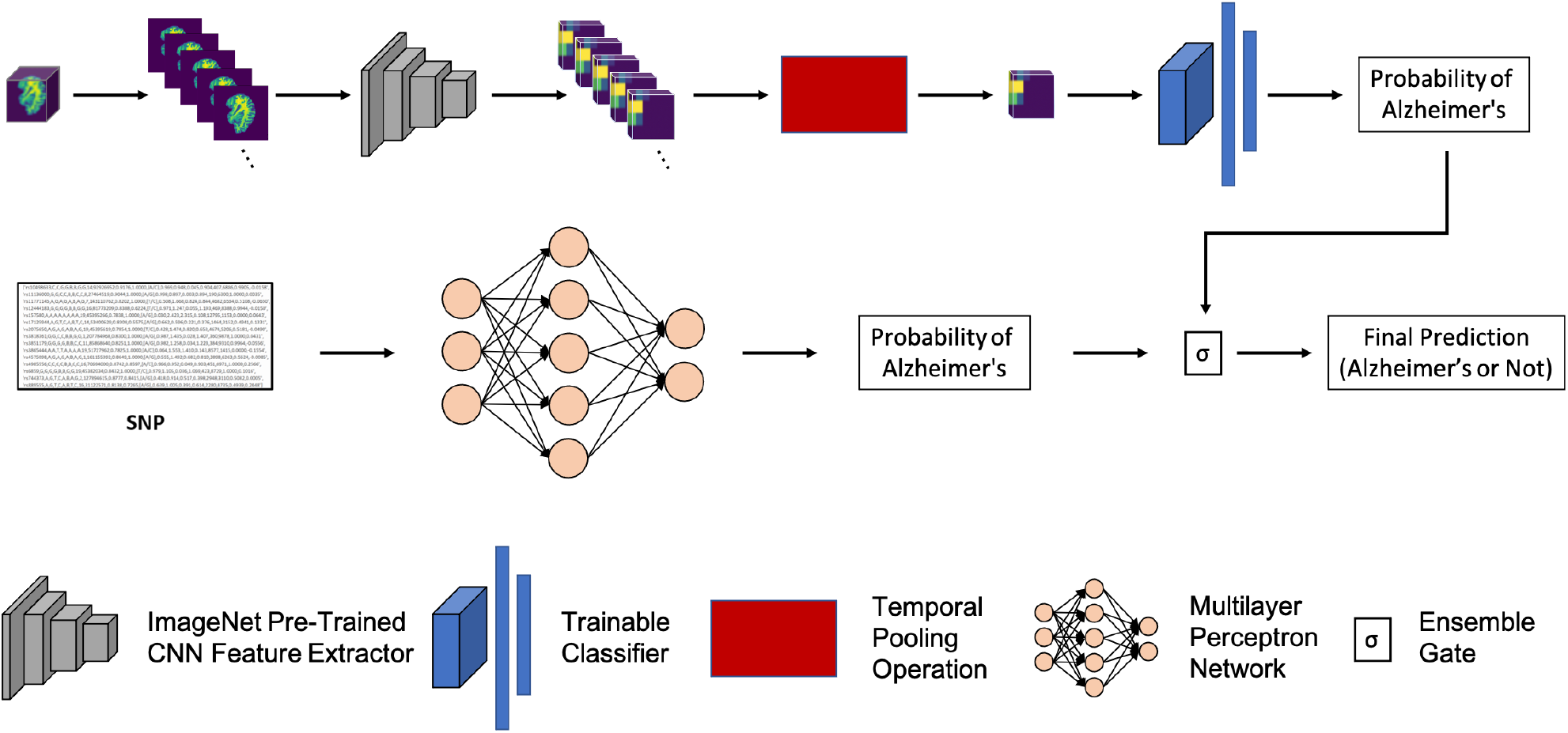
An illustration of the proposed multi-modal model. An MRI and a set of SNPs are feed into a 2D CNN model (top) and an MLP model (bottom), respectively. The ensemble gate opens when the image branch (top) has low prediction confidence, the SNP branch (bottom) is used to refine the overall prediction.

### A. Image Processing Network

The image processing network is a 2D CNN model that is based on our previous work [10]. The 2D CNN model includes three components, a pre-trained feature extractor, a temporal pooling operation, and a shallow CNN classifier. In this work, we adopted the late fusion strategy from [10].

The 2D slices of an MRI are first passed through the feature extract separately. A feature block with the shape of *H* × *W* × *K* × *Z* is extracted, where *W* is the width of a feature map, *H* is the height of a feature map, *K* is the number of feature maps that extracted from one slice, and *Z* is the number of slices in the MRI (*Z* ≥ 1). Temporal pooling operation is then applied to the slice dimension of the feature block. The temporal pooling operation aims to convert the 3D MRI features with different lengths to fixed-size 2D image features by replacing the values on the temporal dimension (or the slice dimension) with a single value. The output shape of the temporal pooling operation is *H* × *W* × *K* × 1. Finally, the fixed-size feature block is used as the input of the shallow CNN classifier for AD diagnosis. Max-pooling is used as the temporal pooling operation according to our previous work. Cross-Entropy loss is used to train the CNN classifier.

### B. SNP Processing Network and SNP Selection

The SNP processing network is a multilayer perceptron (MLP) network that takes a set of SNPs as input and predicts the probability of being the AD class.

In GWAS, millions of common coding and non-coding genetic variants across the genome are tested for association with a trait. Instead of directly genotyped, functional genes usually were found according to linkage disequilibrium (in which restricted recombination between loci causes nonrandom transmission of alleles) with genotyped SNPs. In general, after GWAS identifies hundreds of genetic loci associated with traits, a lot of informatics and functional characterization are still needed to further identify functional variants and genes. For this reason, thousands of SNPs could be identified as associated with one disease phenotype from a single GWAS study, but successfully finding one related functional gene is still challenging.

To effectively utilize SNP data and exploit its application in machine learning, we manually chose 41 SNPs from literature. Due to data availability in the ADNI database, 15 SNPs were used for SNP model construction. Out of these, 8 SNPs have been identified as linked to a functional gene. As shown in Table I, rs11771145, rs17125944, rs10498633, rs2718058 and rs3865444 were chose from a 2013 AD GWAS which used two-stage meta-analysis for 74,046 individuals of European ancestry [21]. In 2018, another GWAS used 314,278 participants from the UK Biobank, with 14,338 CN and 27,696 AD samples [22]. Participants were excluded if their parents were younger than 60 years, died before the age of 60 years, or if no age was reported. In this study, rs12444183, rs4985556 along with their linked genes PLCG2 and IL34 were identified as new risk loci for AD. ADAMTS4 and its lead SNP rs4575098 were chosen from a new GWAS that was published in Nature Genetics, 2019 [23]. This GWAS included a sample size more than eightfold greater than that of the 2013 GWAS by accumulating the genetic data of 635,000 individuals, including a new data set never used before (about 48,000 AD and 330,000 CN).

**TABLE I:** SNPs with Identified Genes

| SNP Name | Chr | Position | Nearest gene | P value |
| --- | --- | --- | --- | --- |
| rs4575098 | 1 | 161155392 | ADAMTS4 | $2.05 \times 10^{-10}$ |
| rs11771145 | 7 | 143110762 | EPHA1 | $8.8 \times 10^{-10}$ |
| rs17125944 | 14 | 53400629 | FERMT2 | $7.9 \times 10^{-9}$ |
| rs10498633 | 14 | 92926952 | SLC24A4 | $1.5 \times 10^{-7}$ |
| rs3865444 | 19 | 51727962 | CD33 | $5.1 \times 10^{-8}$ |
| rs4985556 | 16 | 70694000 | IL34 | $3.7 \times 10^{-8}$ |
| rs12444183 | 16 | 81773209 | PLCG2 | $3.2 \times 10^{-8}$ |
| rs2718058 | 7 | 37841534 | GPR141 | $4.8 \times 10^{-9}$ |

Although no linked functional genes were found for the other 7 SNPs yet, meta-analysis did by [24] using 2429 GWAS studies (which including 1818 phenotypes and 28,462 SNPs) showed that at least two independent GWAS studies identified these SNPs have a significant association with AD (Table II). It is worth noting that nine independent GWAS studies have associated rs2075650 with AD, which is worthy of future functional investigation.

**TABLE II:** SNPs Identified by Multiple Studies

| SNP Name | Chr | Position | Number of independent studies |
| --- | --- | --- | --- |
| rs2075650 | 19 | 50087459 | 9 |
| rs11136000 | 8 | 27520436 | 2 |
| rs157580 | 19 | 50087106 | 3 |
| rs3818361 | 1 | 205851591 | 2 |
| rs3851179 | 11 | 85546288 | 2 |
| rs6859 | 19 | 50073874 | 3 |
| rs744373 | 2 | 127611085 | 3 |

The ADNI study has been divided into several phases,including ADNI-1, ADNI-GO, and ADNI-2, which started in 2004, 2009, and 2011, respectively. Due to the limitation of assessment technology or data collection methods, SNP data for patients from different phases of the ADNI study were not unified. According to our preliminary study, the following values were selected as SNP features for training and testing the SNP model: Allele1-Top, Allele2-Top, Allele1-Forward, Allele2-Forward, Allele1-AB, and Allele2-AB.

### C. Refine the Image Network Performance with SNPs

The SNP processing branch is used to refine the performance of the image processing branch when the prediction confidence is low (i.e., the predicted probability, *p*, is close to 0.5). An ensemble gate, *σ*, opens when *p* is close to 0.5. The SNP processing branch result is assembled with the image processing branch prediction to form the final prediction. A hyper-parameter Θ is used as the threshold to control the ensemble gate that contains two values, *θ_high_* and *θ_low_*. When *θ_low_* ≤ *p* ≤ *θ_high_, σ* is open. Otherwise, *σ* is closed. In order to keep the ensemble stage intuitive and straightforward, we simply average the prediction probability of the two predictions when *σ* is open.

### D. Implementation

The image processing network uses the AlexNet [25] as the backbone. The network is pre-trained on the ImageNet dataset [26]. After the network is trained, the first four convolutional (Conv) layers are used as the pre-trained feature extractor. A 1 × 1 Conv layer and two FC layers with 512 neurons and 2 neurons are added to the feature. The Conv layer aims to convert the ImageNet pre-trained features to AD-specific classification features. The two FC layers are used as an MLP classifier. We implement the network in Pytorch [27] following [10]. Adam [28] optimizer with a learning rate of 10^-4^ is used as the optimizer, and weighted cross-entropy is used as the loss function.

The SNP processing network is an MLP network that is implemented in scikit-learn [29]. The model has 1000 hidden layers with the Relu activation function. SGD with a learning rate of 5 * 10^-3^ is used as the optimizer. The max training iteration is 10,000 steps, but early stopping criteria are applied.

We set *θ_low_* =0.3 and *θ_high_* = 0.7 for the control of the ensemble gate (*σ*). When 0.3 ≤ *p* ≤ 0.7, *σ* opens and SNP prediction is used to refine the image prediction. Otherwise, *σ* remains closed. We use the image prediction as the final prediction of this patient.

## III. Result

### A. Evaluation Setup

T1 MRIs of 100 patients from the ADNI dataset are used in this study, including 51 CN samples and 49 AD samples. The images were pre-processed, the skulls were stripped out from the brain images. Among the 100 patients, 75 patients (35 CN vs. 40 AD) have both image and SNP information. The MRI images were used to train the image processing network. The 5-fold cross-validation is applied to get the prediction result of all the 100 patients. An additional 300 patients (140 CN vs. 160 AD) were selected for the SNP processing network training and validation. The SNPs of the 75 patients who have MRIs are used only for testing the SNP processing network. The image processing network was trained for 100 epochs, and the SNP processing network was trained for 10,000 interactions with early stopping criteria being applied.

The area under the curve of Receiver Operating Characteristics (AUC) and Average Precision (AP) are used as the evaluation metrics. The AUC score considers both the true-positive rate and the false-positive rate. The AP score summaries the area under the precision and recall curve (PRC) that combines precision and recall in a single visualization. For both of the metrics, a higher score indicates a better performance. A classifier that works perfectly would have a score of 1, and a classifier that guessed randomly would result in a score of 0.5.

### B. Classification Performance

Table III shows the evaluation result of i) using only the image processing branch performance (denoted as *Image__X_*, where the subscript, X, indicates the number of patients that was used in the testing), ii) using only the SNP processing branch (denoted as *SNP__X_*), and iii) the proposed method of the 75 patients who have both MRI and SNPs and the overall performance of the total 100 patients. The *Refined__100_* performance of the 100 patients is derived by combining the performance of the *Refined__75_* with the image-only performance of the 25 patients who do not have SNPs. Figure 2 and 3 show the AUC and precision-recall curve (PRC) for the 75 patients and 100 patients, respectively.

**Fig. 2:**
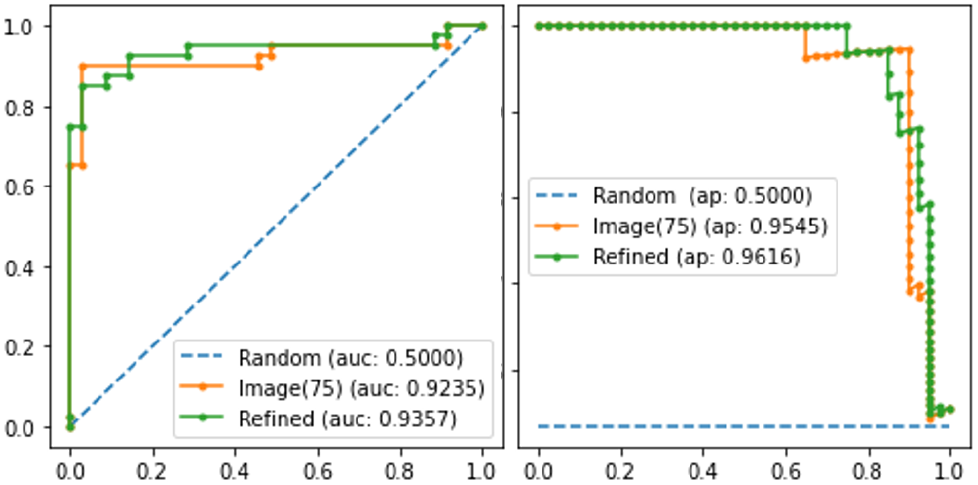
AUC (left) and PRC (right) of the 75 patients who have both the MRI and SNPs. Orange line: Image-only result. Green line: SNP refined result.

**Fig. 3:**
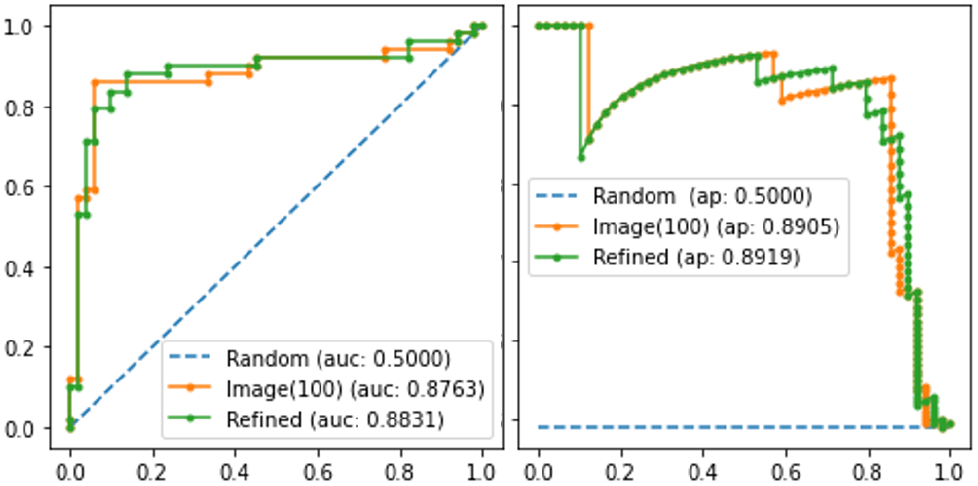
The overall performance of the 100 patients, AUC (left) and PRC (right). Orange line: Image-only result. Green line: SNP refined result.

**TABLE III:** Performance of Different Methods

| Model | # Patients | AUC | AP |
| --- | --- | --- | --- |
| **Image<sub>75</sub>** | 75 | 0.9235 | 0.9545 |
| **SNP<sub>75</sub>** | 75 | 0.6807 | 0.6753 |
| **Refined<sub>75</sub>** | 75 | **0.9357** | **0.9616** |
| **Image<sub>100</sub>** | 100 | 0.8763 | 0.8905 |
| **Refined<sub>100</sub>** | 100 | **0.8831** | **0.8919** |

The result shows that for the 75 patients who have both MRI and SNPs, when using only the image processing network (*Image__75_*), the network gets a 0.9235 AUC and 0.9545 AP. When using the SNPs to refine the image processing network (*Refined__75_*), both the AUC and AP can be improved to 0.9357 AUC and 0.9616 AP. The result reveals that although the performance of the SNP model (*SNP__75_*) was not very impressive (0.6807 AUC and 0.6753 AP), the image model was still beneficial from combining SNP features. Similar results were also obtained in the models using the 100 patients.

## IV. Discussion

### A. 2D CNN vs. 3D CNN

An MRI data is a pseudo-3D image that contains multiple 2D slices. 3D CNNs are a natural choice when applying neural networks to MRI analysis tasks. However, 3D CNNs are usually much larger than 2D models in terms of the number of trainable parameters. Empirical speaking, more trainable parameters usually lead to a longer training time and request more training data, which significantly increases the training cost of 3D CNNs over 2D CNNs. In addition, the small dataset size in this study may also limit the performance of 3D CNN models. Thus, we use the 2D CNN for the image processing branch. According to our previous study [10], the selected 2D CNN model improved the network performance by over 10% compared with the 3D competitor and reduced the overall training time by approximately 95%, from 3,916 minutes to 213 minutes.

### B. Ensemble Gate (σ) Threshold

For a binary classification task, the output of neural networks is often given as the probability (p) of one sample to be a positive class. The possible range of *p* is between 0 and 1, with 0 indicating negative and 1 indicating positive. Normally, the deep learning world may use 0.5 as the cutoff. In our case, any cases with *p* < 0.5 are predicted as CN; otherwise, it is AD. It shows more uncertainty or low confidence in the decision, when p is more close to 0.5.

We propose to use SNPs to refine the image predicting result when the predicted confidence is low. The refinement is done by assembling the image prediction and SNP prediction. An ensemble gate (σ) is used to control the ensemble process. We use 0.3 and 0.7 to define the lower bound and upper bound, respectively, of the low confidence prediction. More specifically, when the imaging predicted, p, within this range (0.3 ≤ *p* ≤ 0.7), we say the prediction has low confidence, σ opens, and SNP prediction is used to refine the image prediction. Otherwise, σ remains closed.

We choose 0.3 and 0.7 as the thresholds partly because it is about one standard deviation from the mean of the range of the possible probabilities. In addition, our SNP model only has about 70% accuracy, which could be inferred as an average 70% prediction confidence or *p* = 0.7 for being positive cases, according to [30]–[32]. Thus, we do not use the SNP to refine the image predictions that have *p* > 0.7 or *p* < 0.3 because the predictions in those two ranges have higher confidence, which hypothetically is more accurate than the SNP predictions.

### C. Limitation of SNP

GWAS have been very successful in identifying novel variant–trait associations [33]–[35]. In the past decades, several large AD GWAS have been published that involved millions of people, which provide us with extensive numbers of SNP information [21]–[23], [36]. Candidate functional SNPs in regulatory and coding regions affect genes differently. Those occurring within the protein coding region can affect protein structure or lead to alternative splicing, potentially resulting in altered function or in some cases loss of function. Genetic variants located in non-coding regions often influence phenotypes by altering the expression of nearby genes [37]. By using this SNP information, 40 risk locus associated with AD have been discovered. Thus, exploiting the application of SNP in AD using machine learning is not only tempting but also promising.

During the experiments, various predictive models and network architectures were evaluated. However, the best result we can achieve is about 70% accuracy, 0.6801 AUC, and 0.6753 AP with an MLP model. Though this number is not very impressive, we think this number is generally acceptable partly because it may be within the range of human experts on many medical tasks. For instance, the average diagnosis performance of radiologists on breast cancer is only 0.71 AUC [38], [39].

One potential limitation for the SNP model performance is limited data availability in the ADNI database. Although hundreds of SNPs were associated with AD, we can only use 15 of them, which are available in the ADNI genetic database. Many newly identified SNPs do not exist in the ADNI database. With the access of more SNPs and a reasonable SNP selection of SNPs, the performance of the SNP predicting model may be improved significantly.

In addition, feature selection may be applied to extract useful features from SNPs. According to ADNI, each SNP has 19 features (Table IV). In the downstream genomic studies of GWAS, Allele1-AB and Allele2-AB are the two commonly used features. However, according to our experiences, these two features cannot provide enough information to the predictive model. We end up using six features: Allele1-Top, Allele2-Top, Allele1-forward, Allele2-Forward, Allele1-AB, and Allele2-AB. In future studies, an optimized feature selection method with statistical and pathological data support might lead to better results.

**TABLE IV:**
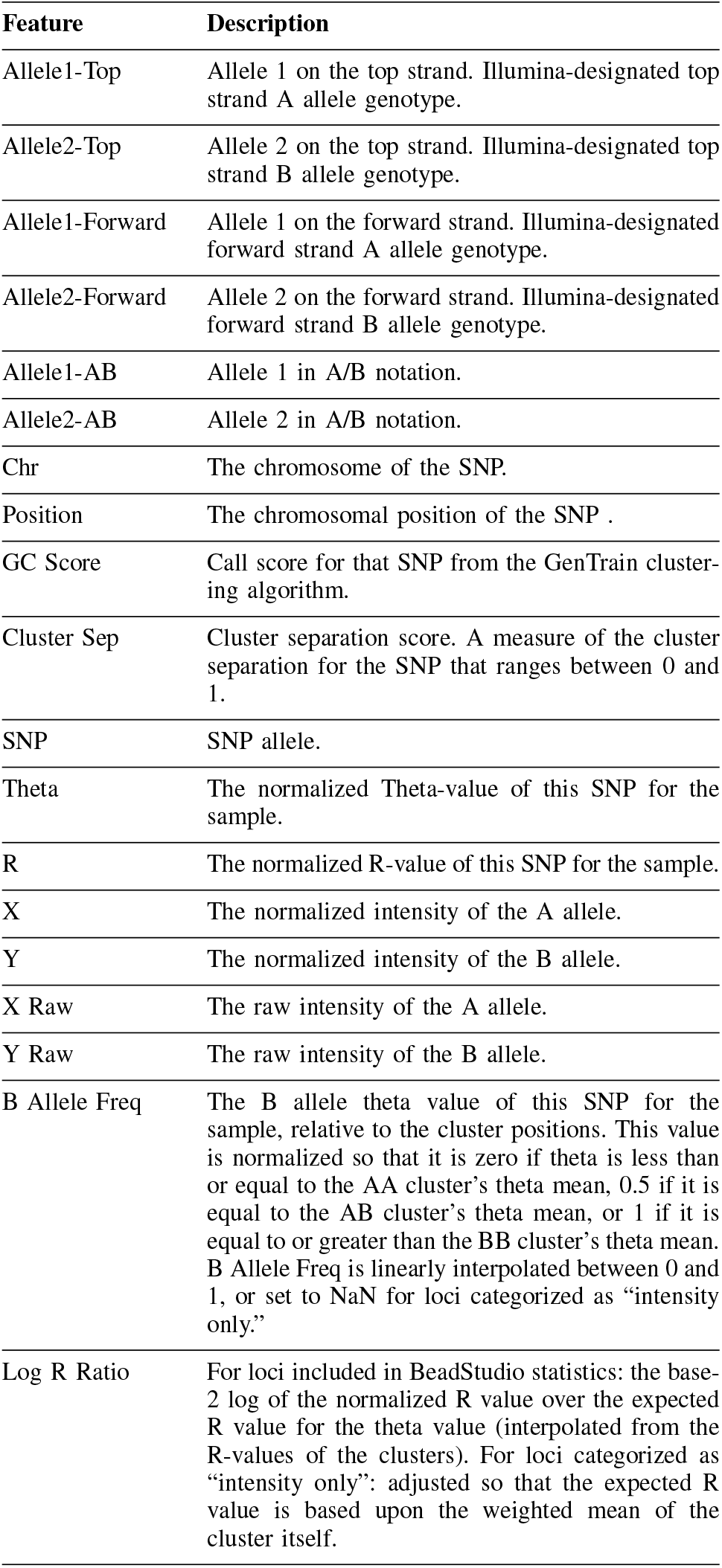
SNP Feature List

| Feature | Description |
| --- | --- |
| Allele1-Top | Allele 1 on the top strand. Illumina-designated top strand A allele genotype. |
| Allele2-Top | Allele 2 on the top strand. Illumina-designated top strand B allele genotype. |
| Allele1-Forward | Allele 1 on the forward strand. Illumina-designated forward strand A allele genotype. |
| Allele2-Forward | Allele 2 on the forward strand. Illumina-designated forward strand B allele genotype. |
| Allele1-AB | Allele 1 in A/B notation. |
| Allele2-AB | Allele 2 in A/B notation. |
| Chr | The chromosome of the SNP. |
| Position | The chromosomal position of the SNP . |
| GC Score | Call score for that SNP from the GenTrain clustering algorithm. |
| Cluster Sep | Cluster separation score. A measure of the cluster separation for the SNP that ranges between 0 and 1. |
| SNP | SNP allele. |
| Theta | The normalized Theta-value of this SNP for the sample. |
| R | The normalized R-value of this SNP for the sample. |
| X | The normalized intensity of the A allele. |
| Y | The normalized intensity of the B allele. |
| X Raw | The raw intensity of the A allele. |
| Y Raw | The raw intensity of the B allele. |
| B Allele Freq | The B allele theta value of this SNP for the sample, relative to the cluster positions. This value is normalized so that it is zero if theta is less than or equal to the AA cluster’s theta mean, 0.5 if it is equal to the AB cluster’s theta mean, or 1 if it is equal to or greater than the BB cluster’s theta mean. B Allele Freq is linearly interpolated between 0 and 1, or set to NaN for loci categorized as “intensity only.” |
| Log R Ratio | For loci included in BeadStudio statistics: the base-2 log of the normalized R value over the expected R value for the theta value (interpolated from the R-values of the clusters). For loci categorized as “intensity only”: adjusted so that the expected R value is based upon the weighted mean of the cluster itself. |

## V. Conclusion

In this study, we propose a novel multi-modal deep neural network. The model uses both MRI and SNPs for Alzheimer’s disease diagnosis. When the imaging branch has low prediction confidence, the SNP branch is used to refine the imaging branch performance. The model is inspired by clinical practice such that when the medical expert is uncertain in his/her decision, additional tests are ordered to help to make the decision. our result shows that the proposed method improves the Alzheimer’s disease diagnosis performance to 93.5% AUC and 96.1% AP, respectively, when all the patients have both MRI and SNP data. We believe this work demonstrates a novel way to use GWAS results in clinical practice. It can serve as a strong baseline for future researchers.

## References

[1] S. Mathotaarachchi et al., “Identifying incipient dementia individuals using machine learning and amyloid imaging,” Neurobiology of aging,vol. 59, pp. 80–90, 2017.

[2] A. Association et al., “2021 alzheimer’s disease facts and figures,” Alzheimers Dement, vol. 17, no. 3, pp. 327–406, 2021.

[3] N. R. Barthélemy et al., “A soluble phosphorylated tau signature links tau, amyloid and the evolution of stages of dominantly inherited alzheimer’s disease,” Nature medicine, vol. 26, no. 3, pp. 398–407, 2020.

[4] R. V. Marinescu et al., “Tadpole challenge: Prediction of longitudinal evolution in alzheimer’s disease,” arXiv preprint arXiv:1805.03909, 2018.

[5] D. Cheng, M. Liu, J. Fu, and Y. Wang, “Classification of mr brain images by combination of multi-cnns for ad diagnosis,” in Ninth International Conference on Digital Image Processing, vol. 10420. International Society for Optics and Photonics, 2017, p. 1042042.

[6] S. Esmaeilzadeh, D. I. Belivanis, K. M. Pohl, and E. Adeli, “End-to-end alzheimer’s disease diagnosis and biomarker identification,” in International Workshop on Machine Learning in Medical Imaging. Springer, 2018, pp. 337–345.

[7] J. Albright, A. D. N. Initiative et al., “Forecasting the progression of alzheimer’s disease using neural networks and a novel preprocessing algorithm,” Alzheimer’s & Dementia: Translational Research & Clinical Interventions, vol. 5, pp. 483–491, 2019.

[8] D. Pan, Y. Huang, A. Zeng, L. Jia et al.,“Early diagnosis of alzheimer’s disease based on deep learning and gwas,” in International Workshop on Human Brain and Artificial Intelligence. Springer, 2019, pp. 52–68.

[9] X. Xing, G. Liang, H. Blanton, M. U. Rafique, C. Wang, A.-L. Lin, and N. Jacobs, “Dynamic image for 3d mri image alzheimer’s disease classification,” in European Conference on Computer Vision. Springer, 2020, pp. 355–364.

[10] G. Liang, X. Xing, L. Liu, Q. Yin, A.-L. Lin, and N. Jacobs, “Alzheimer’s disease classification using 2d convolutional neural networks,” medRxiv, 2021.

[11] Esteva et al., “Dermatologist-level classification of skin cancer with deep neural networks,” Nature, vol. 542, no. 7639, pp. 115–118, 2017.

[12] Q. Yang et al., “Low-dose ct image denoising using a generative adversarial network with wasserstein distance and perceptual loss,” IEEE transactions on medical imaging, vol. 37, no. 6, pp. 1348–1357, 2018.

[13] G. Liang et al.,“Joint 2d-3d breast cancer classification,” in 2019 IEEE International Conference on Bioinformatics and Biomedicine. IEEE, 2019, pp. 692–696.

[14] R. P. Mihail, G. Liang, and N. Jacobs, “Automatic hand skeletal shape estimation from radiographs,” IEEE transactions on nanobioscience, vol. 18, no. 3, pp. 296–305, 2019.

[15] Y. Zhang, X. Wang, H. Blanton, G. Liang, X. Xing, and N. Jacobs, “2d convolutional neural networks for 3d digital breast tomosynthesis classification,” in 2019 IEEE International Conference on Bioinformatics and Biomedicine (BIBM). IEEE, 2019, pp. 1013–1017.

[16] G. Liang, X. Wang, Y. Zhang, and N. Jacobs, “Weakly-supervised self-training for breast cancer localization,” in 2020 42nd Annual International Conference of the IEEE Engineering in Medicine & Biology Society (EMBC). IEEE, 2020, pp. 1124–1127.

[17] W. S. Bush and J. H. Moore, “Genome-wide association studies,” PLoS Comput Biol, vol. 8, no. 12, p. e1002822, 2012.

[18] V. Tam, N. Patel, M. Turcotte, Y. Bossé, G. Paré, and D. Meyre, “Benefits and limitations of genome-wide association studies,” Nature Reviews Genetics, vol. 20, no. 8, pp. 467–484, 2019.

[19] P. G. Ridge et al., “Assessment of the genetic variance of late-onset alzheimer’s disease,” Neurobiology of aging, vol. 41, pp. 200–e13, 2016.

[20] L. Bertram and R. E. Tanzi, “Alzheimer disease risk genes: 29 and counting,” Nature Reviews Neurology, vol. 15, no. 4, pp. 191–192, 2019.

[21] J.-C. Lambert et al., “Meta-analysis of 74,046 individuals identifies 11 new susceptibility loci for alzheimer’s disease,” Nature genetics,vol. 45, no. 12, pp. 1452–1458, 2013.

[22] R. E. Marioni et al., “Gwas on family history of alzheimer’s disease,” Translational psychiatry, vol. 8, no. 1, pp. 1–7, 2018.

[23] I. E. Jansen et al., “Genome-wide meta-analysis identifies new loci and functional pathways influencing alzheimer’s disease risk,” Nature genetics, vol. 51, no. 3, pp. 404–413, 2019.

[24] T. Horwitz, K. Lam, Y. Chen, Y. Xia, and C. Liu, “A decade in psychiatric gwas research,” Molecular psychiatry, vol. 24, no. 3, pp. 378–389, 2019.

[25] A. Krizhevsky, I. Sutskever, and G. E. Hinton, “Imagenet classification with deep convolutional neural networks,” Advances in neural information processing systems, vol. 25, pp. 1097–1105, 2012.

[26] J. Deng, W. Dong, R. Socher, L.-J. Li, K. Li, and L. Fei-Fei, “Imagenet: A large-scale hierarchical image database,” in IEEE conference on computer vision and pattern recognition, 2009.

[27] A. Paszke et al., “Pytorch: An imperative style, high-performance deep learning library,” arXiv preprint arXiv:1912.01703, 2019.

[28] D. P. Kingma and J. Ba, “Adam: A method for stochastic optimization,” arXiv preprint arXiv:1412.6980, 2014.

[29] F. Pedregosa et al., “Scikit-learn: Machine learning in Python,” Journal of Machine Learning Research, vol. 12, pp. 2825–2830, 2011.

[30] C. Guo, G. Pleiss, Y. Sun, and K. Q. Weinberger, “On calibration of modern neural networks,” in International Conference on Machine Learning. PMLR, 2017, pp. 1321–1330.

[31] H. Jiang, B. Kim, M. Y. Guan, and M. R. Gupta, “To trust or not to trust a classifier.” in NeurIPS, 2018, pp. 5546–5557.

[32] G. Liang, Y. Zhang, X. Wang, and N. Jacobs, “Improved trainable calibration method for neural networks on medical imaging classification,” in British Machine Vision Conference, 2020.

[33] A. Sud, B. Kinnersley, and R. S. Houlston, “Genome-wide association studies of cancer: current insights and future perspectives,” Nature Reviews Cancer, vol. 17, no. 11, pp. 692–704, 2017.

[34] L. Duncan et al., “Significant locus and metabolic genetic correlations revealed in genome-wide association study of anorexia nervosa,” American journal of psychiatry, vol. 174, no. 9, pp. 850–858, 2017.

[35] C. L. Hyde et al., “Identification of 15 genetic loci associated with risk of major depression in individuals of european descent,” Nature genetics, vol. 48, no. 9, pp. 1031–1036, 2016.

[36] S. J. Andrews, B. Fulton-Howard, and A. Goate, “Interpretation of risk loci from genome-wide association studies of alzheimer’s disease,” The Lancet Neurology, vol. 19, no. 4, pp. 326–335, 2020.

[37] M. E. Cannon and K. L. Mohlke, “Deciphering the emerging complexities of molecular mechanisms at gwas loci,” The American Journal of Human Genetics, vol. 103, no. 5, pp. 637–653, 2018.

[38] E. B. Cole, Z. Zhang, H. S. Marques, R. Edward Hendrick, M. J. Yaffe, and E. D. Pisano, “Impact of computer-aided detection systems on radiologist accuracy with digital mammography,” American Journal of Roentgenology, vol. 203, no. 4, pp. 909–916, 2014.

[39] X. Wang, G. Liang, Y. Zhang, H. Blanton, Z. Bessinger, and N. Jacobs, “Inconsistent performance of deep learning models on mammogram classification,” Journal of the American College of Radiology, vol. 17, no. 6, pp. 796–803, 2020.

